# Correcting trend bias from preferential sampling in fishery-dependent indices

**DOI:** 10.64898/2026.09.30.755768

**Authors:** Eric J. Ward, Kiva L. Oken, Y. Allen Chen, Kate E. Richerson, Cheryl L. Barnes, Dan S. Holland

## Abstract

Fishery-dependent indices of abundance are often used to inform stock assessments, but fishing effort is generally not randomly distributed. Fishers concentrate sampling in locations expected to be profitable, leading to a form of preferential sampling that can bias model-based indices of abundance. We show that when the strength of preferential sampling is constant over time, it biases only the magnitude of an index, which is absorbed by estimated catchability parameters in most stock assessments. A more interesting case arises when the strength of preferential sampling changes over time; in these cases the index trend can be biased in a way that conventional catchability estimation cannot correct. Using simulations structured on long running trawl survey data from the west coast of the USA for shortspine thornyhead (*Sebastolobus alascanus*), we evaluate two structurally different methods for correcting time-varying trend bias: a joint spatiotemporal model of the density and sampling processes, and a discrete-choice model of fisher location choice. Our simulations include two mechanisms generating preferential biases: effort tracking local density, and effort responding to a spatially structured, but unobserved anomaly that also inflates catch. We show that the ability of the estimation methods to correct bias depends on the mechanism responsible for concentrating effort; while the joint model is robust to both data-generating mechanisms, the discrete choice model is only able to correct biases when the mechanism is anomaly-driven. As a real-world case study, we apply these corrections to simulated preferentially sampled data and include indices in the shortspine thornyhead stock assessment model. We found that uncorrected time-varying preferential sampling positively biased estimates of a major recruitment event and, to a lesser extent, unfished recruitment. This bias propagated through to bias in overfishing limits that grew as the population was projected forward. While bias-corrected indices still produced some positive biases in overfishing limits, these were substantially reduced compared to the naive approach. As the mechanism of preferential sampling is rarely known in advance, our results suggest that the joint model represents a robust default for correcting fishery-dependent indices of abundance.

## Introduction

Abundance indices derived from fishery-independent surveys are a cornerstone of fisheries stock assessment, providing the primary signal of how populations change over time and space.

Increasingly, these indices are produced using model-based approaches instead of design-based estimators, as model-based approaches can accommodate irregular or incomplete sampling, and yield fine-scale predictions of distribution or relative abundance (Shelton et al. 2014; Thorson et al. 2015; Anderson et al. 2025). One of the core assumptions of these approaches is that sampling intensity is independent of the densities of the populations being sampled. Fishery-dependent data on catch per unit effort – which are typically collected in areas with higher average densities – violates this assumption and can lead to biases in indices of abundance. However, fishery-dependent data offer several advantages over fishery-independent data; specifically, fisheries datasets may generate far more observations than scientific surveys, these data are less expensive to collect or are already collected for regulatory purposes, and they may cover seasons or areas that fishery-independent surveys do not sample (Pennino et al. 2019). For this reason, model-based spatiotemporal approaches are increasingly being fit to fishery-dependent catch rates to support stock assessment (Grüss et al. 2023), including cases where fishery-dependent indices are validated using fishery-independent surveys (Cheng et al. 2023).

When sampling effort is concentrated in areas where a species is more (or less, in the case of species avoidance) abundant, the observations are preferentially sampled and spatiotemporal models may return biased indices of abundance (Diggle et al. 2010). When density-driven preferential sampling overrepresents high-density areas (Figure 1), resulting indices of abundance tend to be inflated with biases that may not be consistent over time. These potential biases may not be immediately evident, as naive but flexible spatiotemporal models may fit preferentially sampled data well. The problematic assumption made by these models is that high-density cells are representative of the broader population. Diggle et al. (2010) formalized this preferential sampling concept in the geostatistical literature and illustrated that spatial random effects alone cannot remove the biases. Conn et al. (2017) extended these concepts to ecological data, and showed that jointly modeling data and sampling processes can substantially reduce the biases arising from moderate preferential sampling. Preferential sampling can arise from different mechanisms, however, making the type of correction used important. In the formulation of Diggle et al. (2010) and Conn et al. (2017), sampling effort is density-driven. Alternatively, effort may respond to local conditions that analysts are unable to observe; we refer to this alternate mechanism as anomaly-driven (Figure 1).

**Figure 1.**
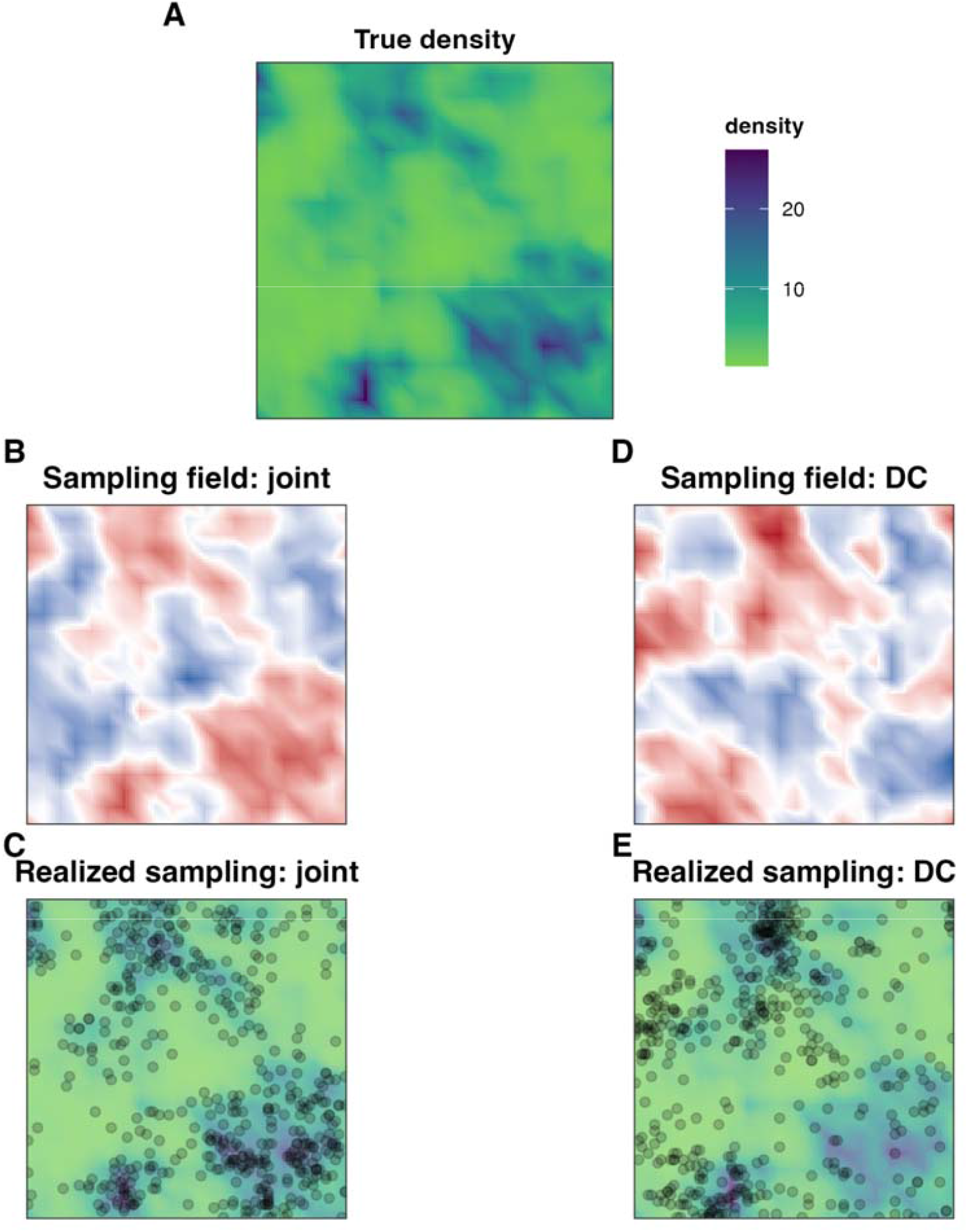
Illustration of the differences between two alternate approaches to preferential sampling. Under the joint model, the probability of a cell being sampled is a function of density (A, B) and fishing effort (black points) is concentrated in high density areas (C). Under discrete choice sampling, fishers’ private information guides the distribution of effort (D) resulting in effort being concentrated in areas that are not always associated with the highest density (E).

Several approaches have been proposed to correct for potential biases associated with preferential sampling. The first approach, developed by Conn et al. (2017) involves a joint model of density with a stochastic sampling process. The sampling process is modeled on a uniform grid, where each cell is modeled as a Bernoulli response with log-odds that are dependent on the underlying expected density field. This formulation builds on the original statistical model of Diggle et al. (2010), and the separation of density and sampling fields developed by Pati et al. (2011). The main assumption of this approach is the form of the density-sampling relationship, which is typically taken to be log-linear. Because the sampling likelihood ties the probability of sampling a cell to the expected density field, this approach is specific to density-driven selection. The likelihood of the joint model draws information about density from cells that are never sampled, which may be inappropriate when cells are not fished for reasons unrelated to abundance (e.g., due to cost, distance from port, regulations, processing capacity; Hoyle et al. 2024).

A second approach to correcting for potential biases associated with preferential sampling is discrete choice (also referred to as random utility) models originating in the fisheries-economics literature. Discrete choice models represent each fishing event as a choice among candidate locations, where the probability of selecting a location increases with its expected utility (Chen et al. 2023; Chen et al. 2026). Utility may include functions of expected catch rates, distance from port, potential risks, and/or costs (Girardin et al. 2017). When information about the drivers of location choice are unknowable to the analyst, fishers may concentrate in locations that appear to have low expected densities under the covariates available to the analyst, but that the fisher knows to be productive (the mechanism being anomaly-rather than density-driven). Like the stochastic sampling model described above, discrete choice models are able to estimate a latent density that may be used to generate an index of abundance. The main difference in these approaches is in their assumptions. Joint models assume stochasticity in sampling with an underlying density field, whereas discrete choice models require assumptions about the form of the utility function and candidate set of alternative locations (Chen et al. 2023; Chen et al. 2026).

An area of research that has received little attention is whether the intensity of preferential sampling in fisheries changes over time, and what effects those changes have on indices of abundance. When the degree of preferential sampling is constant across time, year-specific abundance estimates become inflated (or deflated, in the case of species avoidance) by a similar factor. This type of bias affects the magnitude of estimates, while preserving the trend. Such constant level biases have little impact on stock assessment models, which generally estimate a catchability scaling parameter, especially for fishery-dependent indices (Wilberg et al. 2009).

When the strength of preferential sampling changes over time as a result of changing regulations (such as choke species), costs, or technology, introduced biases affect the estimated magnitude and trend, and cannot be captured with a single catchability parameter in assessment models. For example, the closure of a processing plant may make previously popular high-density fishing areas unprofitable due to the increased travel time to reach markets (Kyle Retherford, pers. comm.). Whether joint or discrete choice models are able to account for time-varying preferential sampling and recover unbiased trends in abundance remains largely unexplored.

Similarly, whether joint or discrete choice models are able to recover indices from data generated under different preferential sampling mechanisms is also unknown.

The objective of this paper is to better understand biases in indices generated from fisheries-dependent data and investigate whether joint spatiotemporal models or discrete choice models can correct for preferential sampling to recover unbiased indices of abundance. We compare a conventional (“naive”) index standardization model that ignores the sampling process to two modeling approaches that include preferential sampling: a joint model that follows the joint sampling model of Conn et al. (2017) and a discrete choice model following Chen et al. (2023). To test whether each correction depends on the mechanism that generated the preferential sampling, we simulate data under both density-driven and anomaly-driven selection and evaluate whether each model recovers the true index under each mechanism. Although previous work has shown that naive spatiotemporal models may be robust to weak and/or time-invariant preferential sampling (Ducharme-Barth et al. 2022), our focus is on whether the joint model can improve index estimation when the degree of preferential sampling changes over time. Using a long-term fishery-independent survey from the West Coast of the USA, we use catches of shortspine thornyhead (*Sebastolobus alascanus*) to generate simulated preferential datasets. In addition to evaluating biases in abundance indices produced from these data, we examine the sensitivity of the current shortspine thornyhead stock assessment model to these indices that are corrected for preferential sampling in different ways.

## Methods

### Simulation design

We used a simulation approach to impose a known sampling process on a realistic latent density surface. To do so, we fit a spatiotemporal model to survey data, treated as known truth, and then used model predictions to generate resampled datasets under varying degrees of preferential sampling. For each simulated dataset, we fit estimation models that either ignored or explicitly accounted for the sampling process. We then compared the estimated index from each model against our known truth and evaluated the sensitivity of recent stock assessments to these estimated indices. All models were implemented in R (R Core Team 2026) and code is provided in our Github repository https://github.com/pfmc-assessments/preferential-simulation and in the sdmTMB R package (Anderson et al. 2025).

### Operating model

#### Simulating fish densities

To generate a realistic latent density surface and trend, we first fit a baseline geostatistical delta-GLMM to West Coast Groundfish Bottom Trawl Survey (WCGBTS, Keller et al. 2017) using catches of shortspine thornyhead (*Sebastolobus alascanus*). We used shortspine thornyhead as a case study species because they are commonly sampled by the WCGBTS (3262 of 6397 hauls), the survey index has shown a marked increasing trend in recent history, and they are a commercially-important species for some segments of the groundfish fishery, with coastwide ex-vessel revenue totaling ∼$2.7 million in 2025. True occurrence rates near 50% allow preferentially sub-sampled simulated data to have a large effect on the occurrence probabilities, and boundary conditions (probabilities near 0 or 1) are less of a concern. Although the WCGBTS time series covers 2003 to present (no survey was conducted in 2020), we subset the data to 2012–2022 so that we could generate shorter 10-year datasets (no data were collected or generated in 2020). Because there have been a number of management changes to different West Coast sectors in the last 20 years (Holland et al. 2025), the duration of consistent fishery-dependent CPUE time series are generally similar to those used here. We fit the baseline index model used in the recent shortspine thornyhead stock assessment (Zahner et al. 2023 and *indexwc* R package, Wetzel et al. 2026) using the R package *sdmTMB* (Anderson et al. 2025).

We generated a triangulated mesh within the WCGBTS survey domain using a cutoff distance of 20 km (points separated by smaller distances are assigned to the same knot or vertex; Rue et al., 2009). In GLMM notation, we can express this index standardization model as

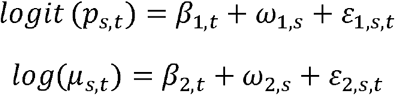

where *p*_*s,t*_is the probability of encountering shortspine in year *t* at location *s, µ*_*s,t*_ is the expected catch conditional on presence, *β*·, _□_ are fixed year effects, *ω*·,_□_ are spatial random fields, and *ε*_□, □_ are spatiotemporal Gaussian random fields; indices 1 and 2 correspond to the encounter (binomial) and positive-catch (Gamma) components. The expected density is the product of the two components, *d*_*s,t*_ = *p*_*s,t*_ *µ*_*s,t*_, and we express normalized log density as *z*_*s,t*_ · Spatial dependence was approximated by the Stochastic Partial Differential Equation (SPDE) and Gaussian Markov random field method (Lindgren et al. 2011). The fitted model was then used to make predictions onto a uniform 2 km by 2km grid within the survey domain for each year.

These predicted density surfaces served as known truth from which all observations were simulated and against which all estimated indices were compared.

#### Simulating preferential datasets

To simulate preferential data under the joint sampling model, we first simulated a sampling intensity field that allowed certain types of habitats (e.g., higher density, closer to ports) to have a higher probability of sampling. We simulated a smooth Gaussian random field for sampling intensity (*η*) shared across years (Matérn range 200 km, marginal standard deviation = 1). Then the probability that grid cell *s* was sampled in time *t* can be expressed as:

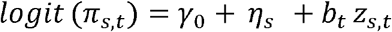

where *γ*_0_ is a coefficient (held at -2) that sets the baseline sampling rate, *η*_*s*_ is a spatial field that represents sampling intensity and is independent of density (e.g., effect of distance to port, etc), *z*_*s,t*_ is the standardized log of true expected density, and the coefficient *b*_*t*_ reflects the strength of density-driven preferential sampling. When *b*_*t*_ = 0, sampling is independent of density. When *b*_*t*_ > 0, sampling is concentrated in high density cells. We explored average values of *b*_*t*_ between 0 and 5. A change in z-scored log density of +1SD from the mean increases the sampling probability for an average cell from 0.12 to 0.27 when □ = 1, 0.12 to 0.73 when □ = 3, and 0.12 to 0.94 when b = 5. While Conn et al. (2017) applied *b*_*t*_ to spatial and spatiotemporal random field anomalies rather than overall expected density (which includes fixed effects), our simulation model is consistent with this because the scaled density has mean 0, ensuring *b*_*t*_ isolates relative density anomalies without confounding annual fixed intercepts. Our joint estimation model, described below, is consistent with the original formulation of Conn et al. (2017).

To simulate data under the discrete choice (anomaly-driven) sampling model we replaced the density-driven mechanism above with structured catch anomalies as predictors. We simulated a smooth Gaussian random field shared across years (Matérn range 40 km, marginal standard deviation = 1) and treated it as the anomaly, or private fisher information, that is unobserved (representing, for example, knowledge of ephemeral fish behavior impacting catchability). In each year, we modified the probability of cells being sampled to be proportional to the anomaly 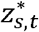 (instead of log density *z*_*s,t*_),

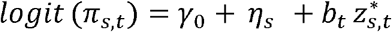

For each simulated dataset, we sampled 1,000 locations without replacement for each year (1, …, 10), with selection probabilities proportional to *π*_*s,t*_. At each sampled location, we simulated a delta-lognormal observation from the true field, modeling encounter probability *p*_*s,t*_ as a Bernoulli response and positive catch from a lognormal distribution. Bernoulli probabilities were generated from the spatiotemporal model predictions fit to the density data. For the density-driven model, we generated catches as a function of the predicted catch rate from the spatiotemporal model predictions,

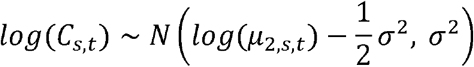

where the log-scale standard deviation 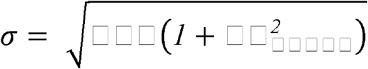, and 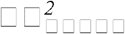 is the coefficient of variation estimated in the delta-gamma model fit to the shortspine density data. For the anomaly-driven model, we adjusted expected catches to also be influenced by the anomaly 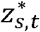,

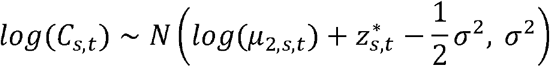

For each simulation model (density-driven, anomaly-driven), we considered two regimes for the strength of preferential sampling *b*_*t*_: constant and time-varying. For the constant regime, we held *b*_*t*_ constant across years at values of 0, 3 (intermediate preferential sampling), or 5 (strong preferential sampling). For the time-varying regime, we assumed *b*_*t*_ to increase with a linear trend from X up to *b*^*^, plus stationary AR(1) deviations (*p*;= 0.7, sd = 0.3), using the same values of *b*^*^ as in the constant scenario. Our primary focus was on time-varying preferential sampling because it has the greatest potential to distort the estimated index. Because the preferential sampling field and the true population trend were identical across replicates, variation in each scenario (n = 25) reflected only stochastic sampling locations and observations.

### Estimation models

We fit three estimation models to each simulated data set. All three used a delta-lognormal observation model with SPDE-based spatial and spatiotemporal Gaussian random fields, and included year-specific fixed intercepts. A simplifying assumption of the estimation models compared to the simulation models was that the spatial and spatiotemporal range was shared as a single parameter, rather than estimated separately. Consistent with other implementations of the SPDE approach, we placed weakly informative penalties on the variance and range parameters (Table in SI). Our first model, representing a naive model with no bias correction, consisted only of the observation likelihood, similar to spatiotemporal models that are fit to fishery-independent data.

Our second estimation model represents the joint model, following the formulation of Conn et al. (2017), extending the naive model with an explicit sub-model for the sampling process. We assumed the sampling process was the same as in the simulation model, where the log-odds that a cell is sampled in a given year depends on that cell’s expected log density *z*_*s,t*_ and a latent field *η*_*s,t*_. Sampling was represented as a Bernoulli response over every cell of the survey grid, so the component spatial deviations (consisting of spatial and spatiotemporal fields, *δ*_*s,t*_ =*ω*_*s*_ +*ε*_*s,t*_), inhomogeneous process is represented exactly rather than approximated. We generated positive- and related these to the indicator *R*_*s,t*_ that cell is sampled in year □ over the full grid,

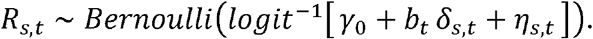

For the time-varying *b*_*t*_ estimation model, we allowed the strength of the density-dependent sampling effect, *b*_*t*_, to vary through time, but we distinguished the estimation model from the simulation model by specifying *b*_*t*_ as a random walk.

Our third estimation model is an extension of the discrete-choice model described by (Chen et al. 2023) where each observation is treated as a choice among a set of candidate cells. For observation □ with chosen cell □ _□_ in year □ _□_ and candidate set □ _□_, the probability that □ _□_ is chosen over candidate cells is,

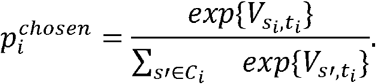

The utility of a candidate cell combines its expected log density with a latent selection field,

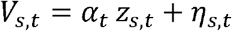

where *α*_*t*_ is a year-specific preferential-strength coefficient (a random walk across years, similar to *b*_*t*_ in the joint model) and *η*_*s,t*_ is a latent Gaussian random field, shared across years, representing density-independent structure in selection. In many economics models, *η*_*s,t*_ is not included because other proxies are available (expected revenue, distance to port), but because those components are not included in our simulation we use *η*_*s,t*_, as in the joint model. For each observation, we constructed a set of 25 total points, randomly sampling cells within the 4 closest SPDE mesh vertices (nodes). Following the control-function approach, the positive-catch predictor is adjusted with a correction (Chen et al. 2023). Though this is often implemented with fixed effects, we treat the coefficients as spatially varying second-order polynomials in the choice probability, together with the latent field,

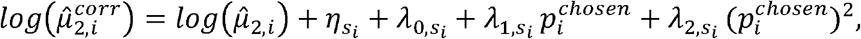

Where *λ*_0_ *λ*_1_ *λ*_2_ are spatially varying coefficients (each a Gaussian random field). The fields 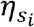 and 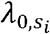 are expected to be highly correlated, but not perfectly so, as only *η*_*s,t*_ appears in the utility model. For simplicity, we share the range of the three spatially varying coefficients as a fixed parameter, but allow them to have separate variances. The latent field *η*_□, □_ thus enters both the utility and the catch predictor and is identified primarily from the catch anomaly.

### Index standardization and evaluation

From each fitted model, we computed an annual abundance index as the area-weighted sum over grid cells of predicted density (estimated density × cell area), ignoring latent effects *η*_*s,t*_. Indices were evaluated on two prediction grids: one that spans the entire survey area and one that includes a 10-km buffer around simulated sampling locations that reflects the shrinking spatial footprint under preferential sampling. We summarized performance across replicates and models by scaling each estimated index relative to its first year to quantify relative trends.

### Impact on stock assessment models

We used simulated indices as inputs into the most recent stock assessment model for shortspine thornyhead (Zahner et al., 2023) to understand how preferential sampling biases may impact estimates of key population parameters. The assessment model for shortspine thornyhead is a sex-specific age-structured integrated population dynamics model that is fit to data from 1901 to 2022. The model is built using Stock Synthesis (Methot and Wetzel, 2013). It has three fishing fleets, all with catch data, discard rates, and length compositions for retained and discarded fish, and three survey fleets (including the WCGBTS) with length compositions and fishery-independent indices. The shortspine thornyhead assessment model does not include age data because there is no validated ageing method. As such, natural mortality and growth parameters are fixed based on a small research dataset (assumed M = 0.04 yr^-1^; Zahner et al., 2023).

We added the simulated indices (2012 to 2022) to the stock assessment model as a fourth survey fleet. For simplicity, we assumed that selectivity was identical to that of the WCGBTS that was used to simulate data for the new indices (i.e., length compositions were unaffected by preferential sampling). We assumed a linear relationship between the index and selected biomass with a lognormal error distribution. Index data are weighted in stock assessments by their standard errors (smaller standard errors translate to more weight); input log standard errors were estimated by the index standardization models. We did not allow the assessment to downweight the index through an additional standard error. If preferential sampling is adequately accounted for, this process should upweight the WCGBTS index from 2012-2022.

We compared assessment model estimates of unfished recruitment (R_0_), the 2008 log-recruitment deviation (ε_2008_), and the 2025 and 2034 overfishing limits (OFL_2025_ and OFL_2034_) across two index standardization methods (naive, joint models) and three levels of preferential sampling (b = 0, 3, 5). In the most recent stock assessment model, 2008 had the second-highest recruitment deviation of the entire time series. This high recruitment estimate is partially driven by a generally monotonic increase in the WCGBTS index during the years used to generate simulated indices. Therefore, we expected that the magnitude of the 2008 recruitment deviation would be sensitive to preferential sampling and how it was accounted for. For catch advice, 2025 is the first year the assessment was used for management and 2034 is the final year of the projection period. The “true” value of all four of these quantities is unknown because, aside from the new index, the underlying data are not simulated. Instead, we considered the naive standardization model with no preferential sampling (b = 0) as the baseline for comparison as it allowed us to quantify biases introduced by preferential sampling. We did not include estimates from the discrete choice model because our primary interest is in how any trend bias affects stock assessment outputs, regardless of the estimation method used to generate the index.

## Results

For each combination of estimation model (joint sampling model, discrete choice model), data-generating process (density-driven, anomaly-driven), and preferential-sampling strength, we computed the Root Mean Squared Error (RMSE) and log-linear trend slope to compare estimated biomass trajectories to truth. In our simulation scenario when the strength of preferential sampling was static in time, we found that greater preferential sampling affected the level (offset), but not the trend (Figure 2). As this offset is generally absorbed in catchability within stock assessment models, we focus the majority of our interpretation on the time-varying preferential sampling scenarios.

**Figure 2.**
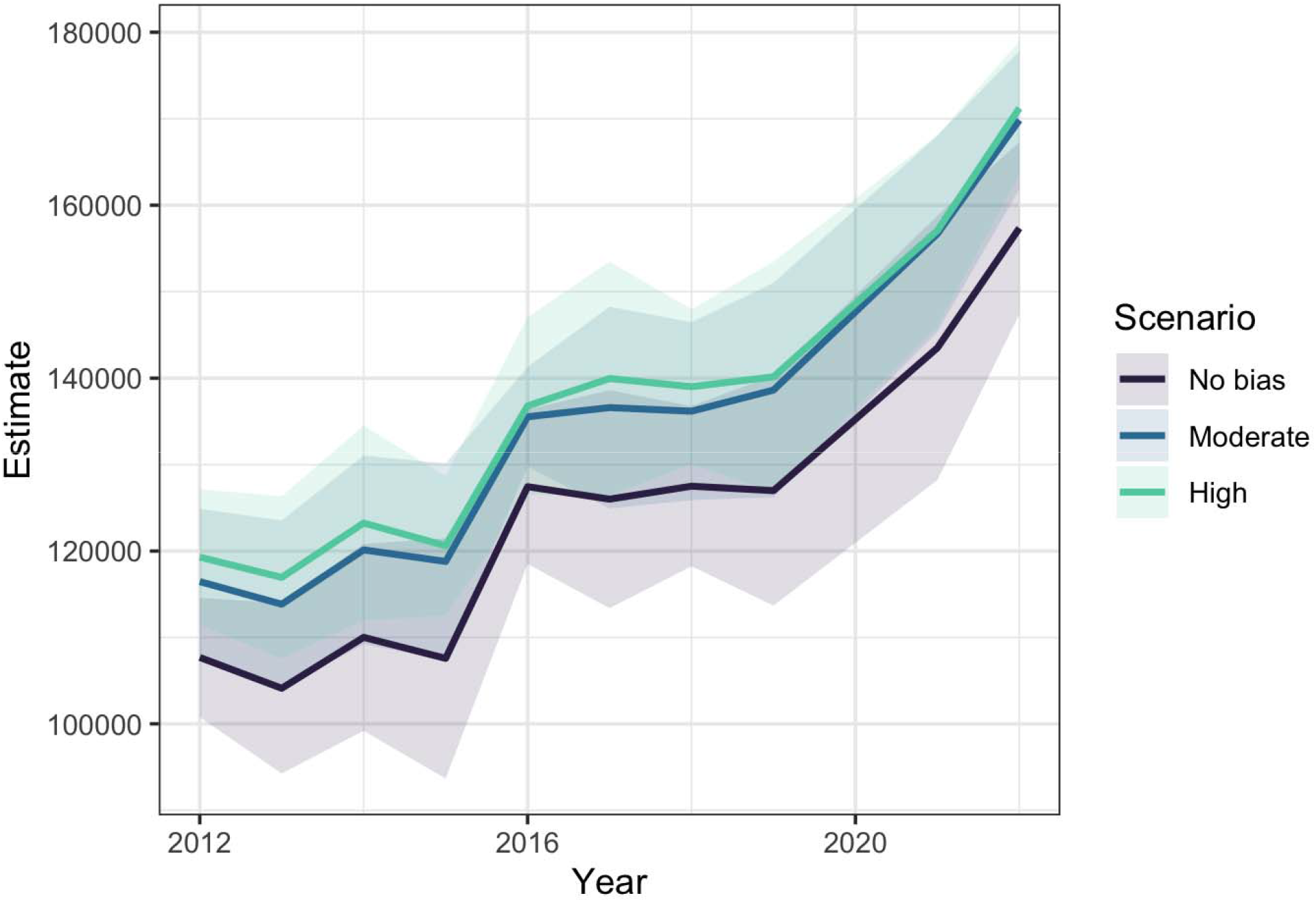
Estimated indices under density-driven preferential sampling scenarios. Scenario levels correspond to preferential coefficients of 0 (‘No bias’), 3 (‘Moderate’) and 5 (‘High’) respectively. For each scenario, 10 datasets were simulated and fit using the software sdmTMB (Anderson et al. 2025); ribbons represent the 95% confidence intervals by scenario and year, and solid lines represent the mean.

### Density-driven sampling

Under the density-driven data-generating process, the naive estimator was positively biased, overstating the index trend by an average of 9% inflation of the true slope (Table S2; Figures 3-4). This bias also increased as the strength of preferential sampling increased. The joint model substantially corrected this bias, removing ∼ 75% of the naive bias under moderate preferential sampling (b = 3) and closer to 90% under stronger preferential sampling (b = 5). In contrast, the discrete choice estimation model did not correct the bias and instead inflated the slopes (more so than the naive estimation model with no bias correction applied; Table S2; Figure 4).

**Figure 3.**
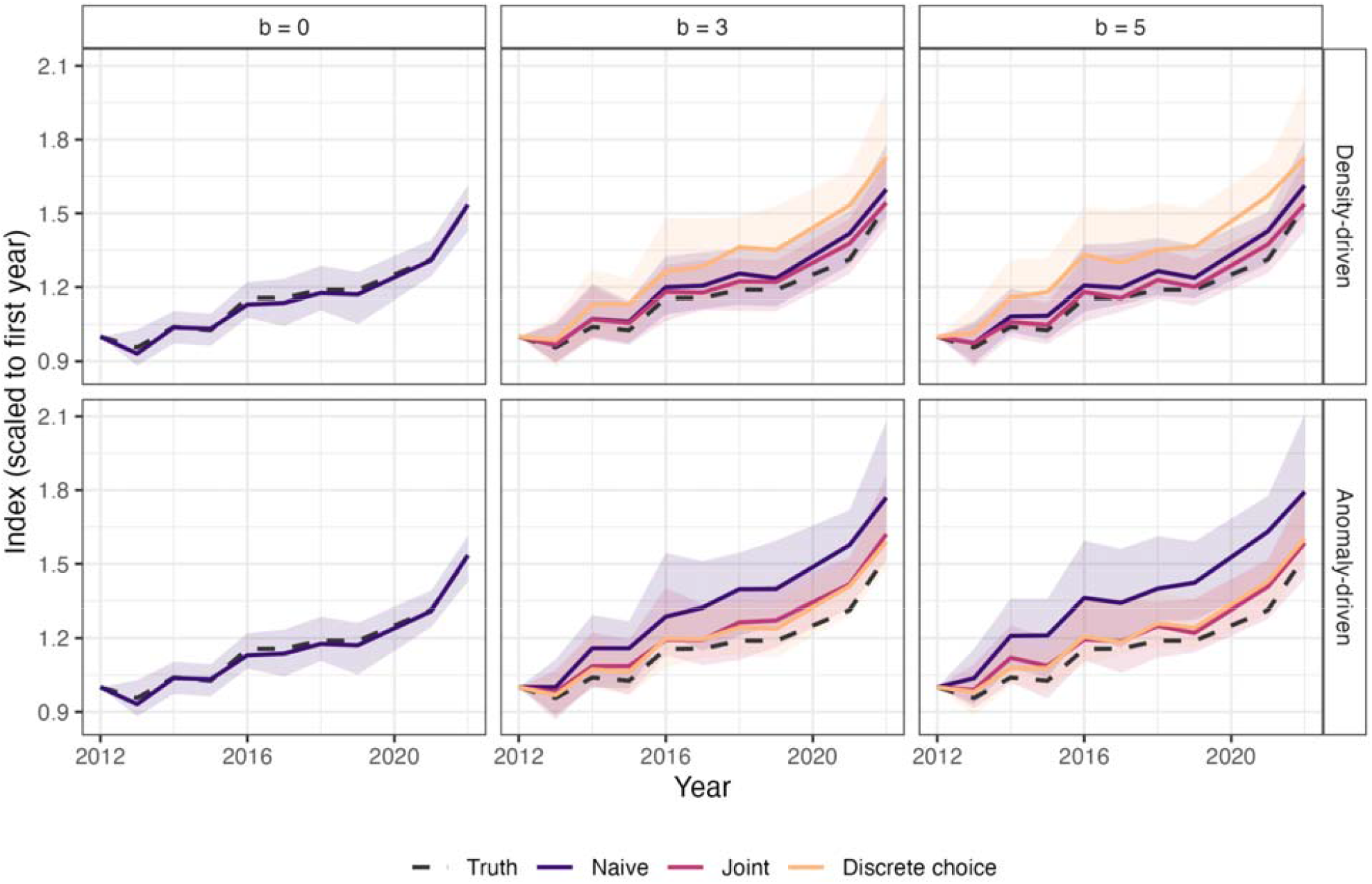
Results from our simulated datasets, summarized across replicates for data generated from the density-driven and anomaly-driven processes. For each level of preferential sampling (b = 0, 3, 5), we show the median and 95% confidence intervals for trend estimates from the naive (no preferential sampling), joint, and discrete choice models. All biomass indices are scaled to the first year (1.0); preferential sampling increases with increasing b and over time, leading to increased biases with the naive model.

**Figure 4.**
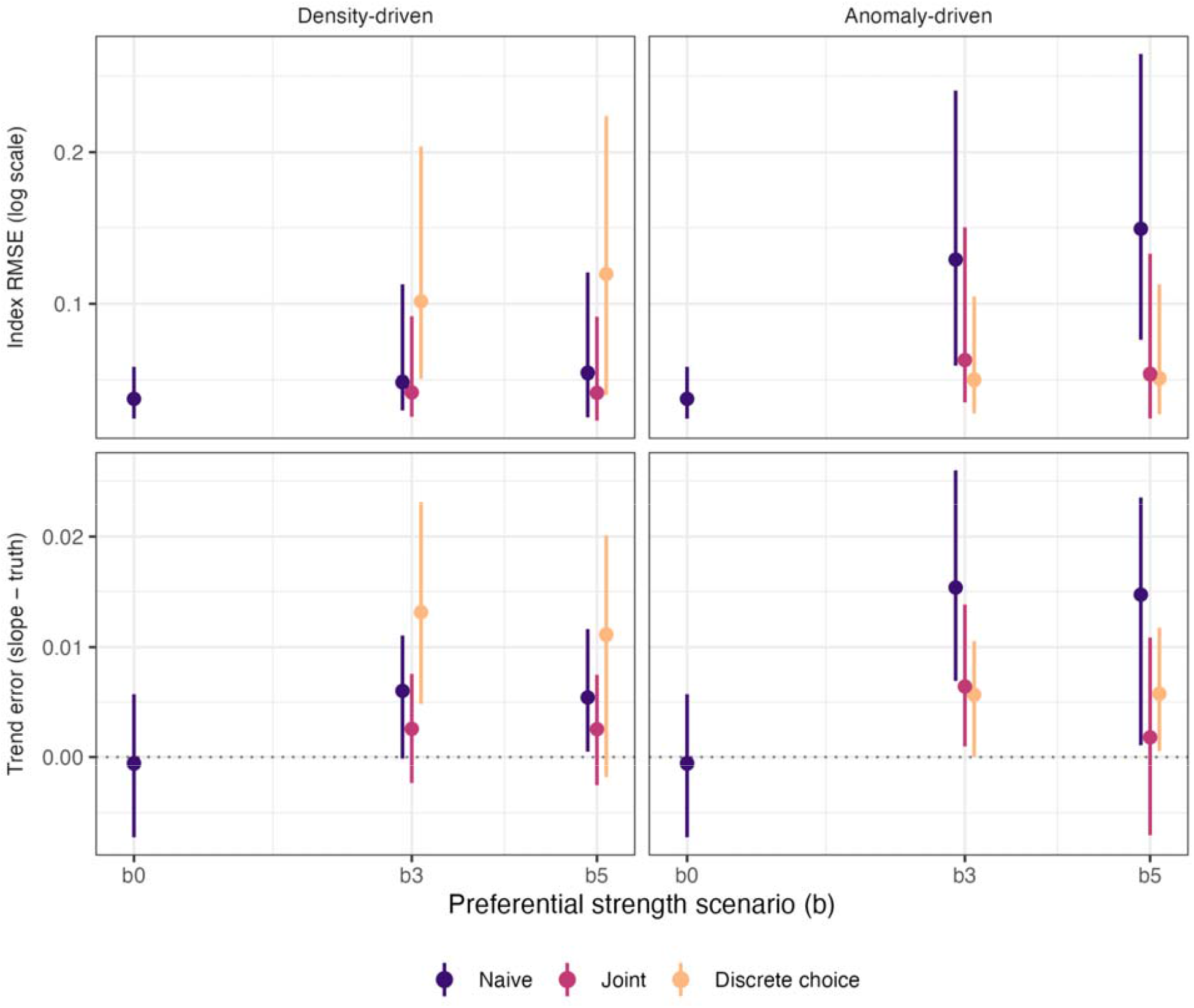
Summaries of estimated biomass indices by estimation model (colors) and preferential sampling mechanisms (columns). The top row shows Root Mean Squared Error (RMSE) between the estimated biomass index from each model (Naive, Joint, Discrete Choice) and the true index; lower values are better. The bottom row shows the bias in the estimated trend slope. For both rows, points represent means across 25 replicates, and vertical bars represent the 95% CIs.

### Anomaly-driven sampling

Under the anomaly-driven process, in which sampling responds to a spatially structured, unobserved catch anomaly that also inflates catch, the naive estimator was more severely biased (overstating the trend estimate by 30%; Figure 4). A major difference between the anomaly-driven simulations and density-driven simulations was that both methods appeared to correct for anomaly-driven preferential sampling. The joint model removed 66% of the bias at b = 3, and 99% at b = 5, while the discrete choice model removed 75% and 64% (Table S2; Figure 4).

### Mechanism of discrete-choice correction

Examining the model fits from the discrete choice models, the correction was driven almost entirely by the latent spatial field rather than the choice-probability polynomial. Across replicates and data generating models, the polynomial coefficients remained at or near initial values, while the latent field standard deviation was consistently identified (Figure S1). Under the anomaly-driven process, inclusion of the latent field recovers the unobserved anomaly and corrects the index. However, when the mechanism is density-driven, this same latent field may partially absorb spatial patterning in density, leading to some of the over-correction described above.

### Impact on assessment models

After we updated the most recent shortspine thornyhead stock assessment model to include our simulated fishery-dependent CPUE indices (the joint and naive models, with and without bias correction, respectively), we summarized quantities of interest across 25 replicates (Table S3). These summaries indicated that estimates of unfished recruitment (R_0_) were positively biased as the strength of preferential sampling increased (Figure 5) but less biased when corrected with the joint model. The 2008 log-recruitment deviation (ε_2008_) appeared to be positively biased with both the naive and joint model, but the bias was larger and affected by the strength of preferential sampling when indices were not corrected for preferential biases (with b = 5, the average uncorrected value is 1.17 and average corrected value 1.08; Figure 5). Estimates of 2025 and 2034 overfishing limits (OFL_2025_ and OFL_2034_) generally appeared to be bias-corrected with the joint model, however the naive model had a positive bias that increased with the strength of preferential sampling, and became worse as the population was projected from 2025 to 2034 (mean OFL_2034_ goes from 1140 to 1160, Figure 5).

**Figure 5.**
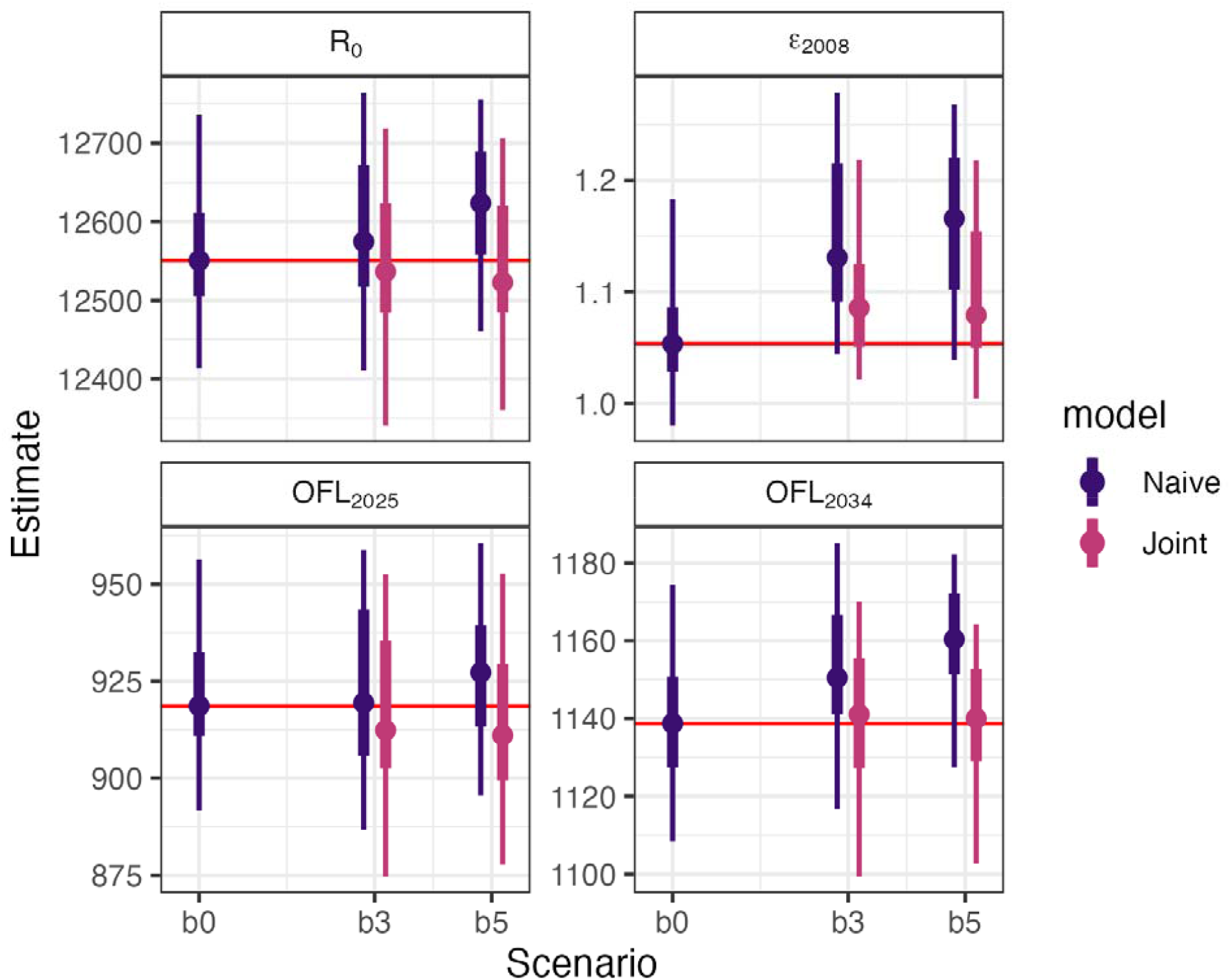
Estimates of key parameters and derived quantities from the shortspine thornyhead stock assessment model after inclusion of indices resulting from naive and joint models: unfished recruitment (R_0_), the 2008 log-recruitment deviation (ε_2008_), and the 2025 and 2034 overfishing limits (OFL_2025_ and OFL_2034_). Points are medians from the simulations, thick lines are 50% quantiles, and thin lines are 95% quantiles. If the assessment model is robust to a given level of preferential sampling and index standardization method, the resulting distributions will look similar to the scenario at b = 0 (no preferential sampling, red horizontal line at median).

## Discussion

Fishery-dependent data are an increasingly attractive source of information for stock assessment (Maunder and Punt 2004), but their value depends on whether biases introduced by non-random samples can be identified and corrected. We show that when the strength of preferential sampling is static over time and estimation methods don’t account for preferential biases, only the level or scale of resulting indices is affected (Wilberg et al. 2009). As most stock assessment models estimate catchability parameters for fishery-independent or fishery-dependent indices, any level biases from fishery-dependent indices are ultimately absorbed by these parameters.

When the strength of preferential sampling changes over time, however, both the level and trend of the index are affected. Using simulations based on trawl survey data from the West Coast of the USA, we show whether and how this bias can be corrected depending on the mechanism of preferential sampling and the correction method applied.

Across both data-generating mechanisms, the joint sampling model of Conn et al. (2017) reduced the trend bias substantially, whereas the discrete-choice model’s ability to correct was influenced by the data-generating mechanism. These findings support the recommendation by Ducharme-Barth et al. (2022) to evaluate joint models under non-stationary dynamics. When the data-generating model was anomaly-driven, with catch anomalies affecting the distribution of effort as well as catches, the discrete-choice model performed as well as or better than the joint model. Under the density-driven data-generating model, however, the discrete-choice model performed consistently worse, inflating the trend beyond that of the uncorrected naive index. The discrete choice model’s strength may not be as a general purpose correction, but as a focused tool for analyzing profit-maximizing behaviors with private information (Chen et al. 2023; Chen et al. 2026).

A practical challenge for stock assessment scientists is that the mechanism of preferential sampling is rarely known in advance. Density-driven and anomaly-driven selection are not mutually exclusive, and some fisheries may exhibit both. In some cases, fishing effort may concentrate in highly productive areas while fishers respond to private information about local conditions (Girardin et al. 2017; Hoyle et al. 2024). Because the joint model in our simulations appears to correct biases under both mechanisms, it may be a more robust choice when underlying mechanisms are unknown. In settings where there is reason to believe that sampling responds to unobserved catch-correlated productivity and potential forms of the utility function are known, the discrete choice model may be a better alternative (providing both the index correction, and mechanistic explanation).

There are several important limitations of our analyses. Though we allowed the strength of preferential sampling to vary over time, the spatial patterning of the latent driver (*η*_*s*_ in our simulations) was constant over time. Real fisheries may experience year to year changes in costs, processing capabilities, or regulations (Hoyle et al. 2024) that may make such assumptions unrealistic. Relatedly, our simulations also use a fixed spatial range for the sampling process (*η*_*s*_), which can be seen as controlling the patchiness of fishing intensity. Future extensions of our modeling framework could investigate the ability to correct for biases when the patchiness changes through time (relative to density, or the spatial scale of utility in the discrete-choice modelling framework). A third limitation is that we made no direct assumptions about schooling or shoaling behaviors (Harley et al. 2001). Although shortspine thornyhead are generally sedentary, such behavior could impact indices developed from preferentially sampled data for other species that do school or shoal, where local densities may become decoupled from broad scale abundance.

Our findings examining the effect of using fishery-dependent indices within the stock assessment model of shortspine thornyhead illustrates one of several use cases. In our assessment modeling framework, we only considered the bias-corrected simulated fishery-dependent CPUE indices alongside fishery-independent trawl survey indices. As a best case scenario, a fishery-dependent CPUE index with biases removed tracks the fishery-independent index, providing more weight to the common population trend. When biases persist in the fishery-dependent index, however, inclusion of fishery-dependent indices may pull the stock assessment away from the survey signal. A second use case involves using fishery-dependent CPUE data in assessments when fishery-independent information is unavailable; in these cases, the entire trend in the assessment model may be driven by the fishery-dependent CPUE index. In these scenarios, bias correction is essential, as all downstream quantities (catch limits, etc) will also be biased.

## Acknowledgements

We extend our thanks to the West Coast Groundfish Bottom Trawl Survey team; without their collection of high quality fisheries independent data, our work would not be possible. We also thank Megsie Siple, whose comments helped us improve the clarity and results of this work.

## Supplemental Information

**Table S1.**
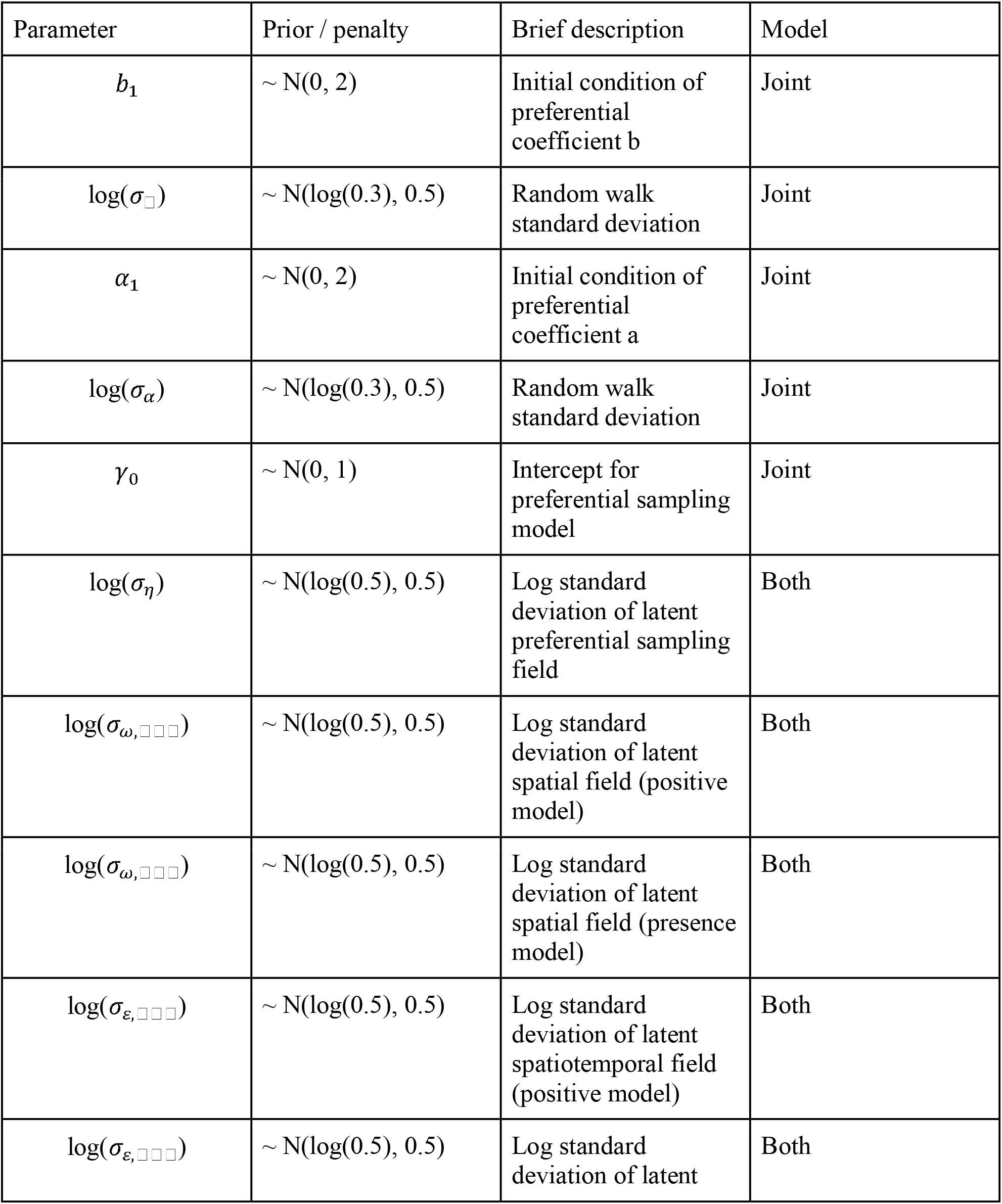
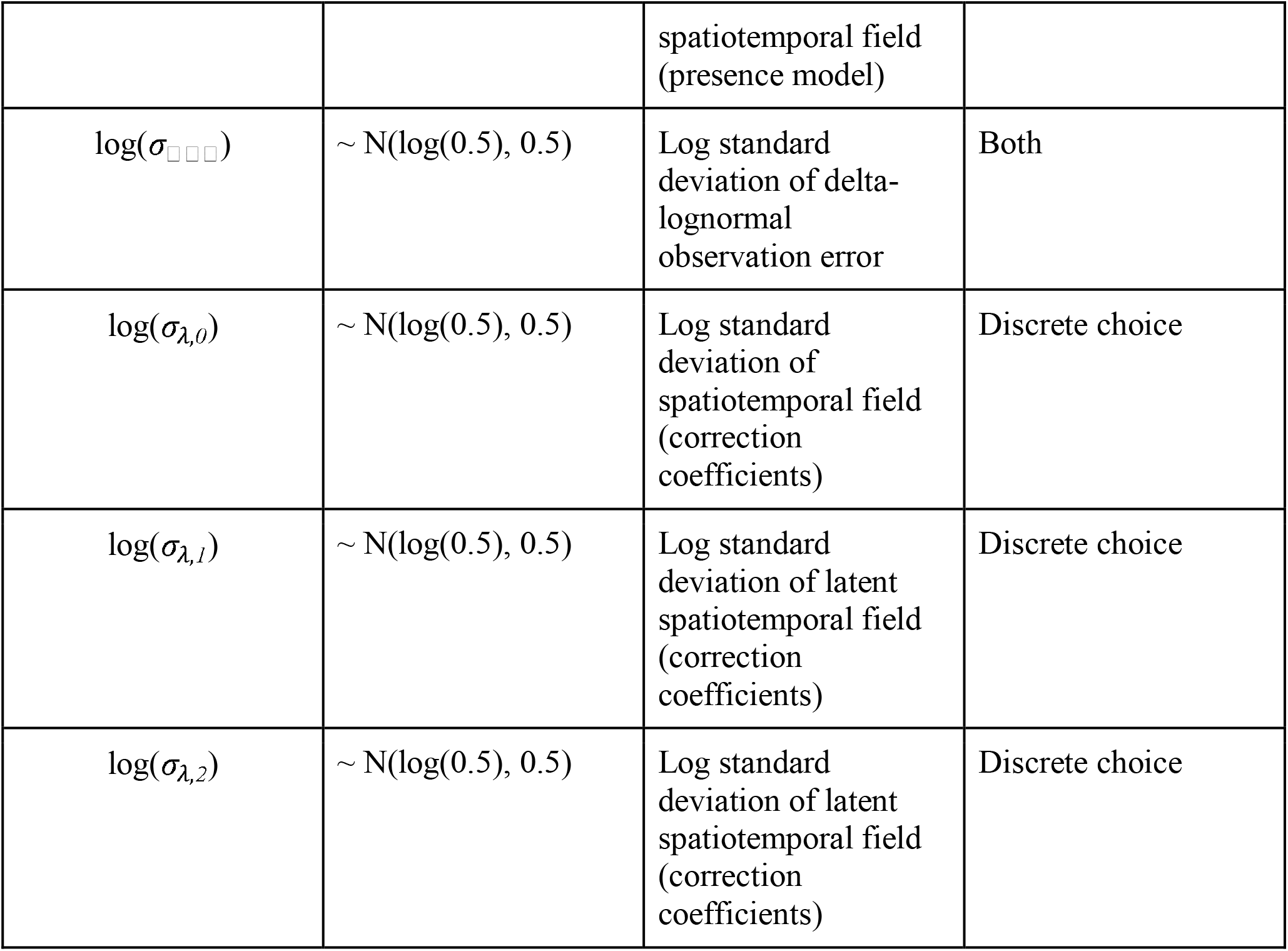
Summary of prior (penalty) values used in our maximum likelihood estimation models.

**Table S2.** Summary of average trend, trend bias and RMSE values calculated across mechanisms, models, and scenarios; n represents the number of replicates (of 25) that successfully converged.

| Mechanism | Scenario | Model | n | E[slope] | Bias | RMSE |
| --- | --- | --- | --- | --- | --- | --- |
| density | b0 | naive | 25 | 0.04 | -1.79E-03 | 0.00408 |
| density | b3 | naive | 25 | 0.0455 | 3.63E-03 | 0.00494 |
| density | b3 | joint | 23 | 0.0427 | 8.69E-04 | 0.00307 |
| density | b3 | dc | 25 | 0.0534 | 1.16E-02 | 0.0127 |
| density | b5 | naive | 25 | 0.0458 | 4.00E-03 | 0.0051 |
| density | b5 | joint | 22 | 0.0423 | 4.59E-04 | 0.00308 |
| density | b5 | dc | 25 | 0.0505 | 8.73E-03 | 0.01076 |
| anomaly | b0 | naive | 25 | 0.04 | -1.79E-03 | 0.00408 |
| anomaly | b3 | naive | 25 | 0.0555 | 1.37E-02 | 0.01465 |
| anomaly | b3 | joint | 21 | 0.0465 | 4.66E-03 | 0.00598 |
| anomaly | b3 | dc | 25 | 0.0453 | 3.47E-03 | 0.00474 |
| anomaly | b5 | naive | 25 | 0.0535 | 1.17E-02 | 0.01332 |
| anomaly | b5 | joint | 19 | 0.0419 | 8.55E-05 | 0.00487 |
| anomaly | b5 | dc | 25 | 0.046 | 4.19E-03 | 0.00529 |

**Table S3.** Summary statistics for shortspine thornyhead assessment model output (data presented in Figure 5).

| scenario | model | metric | median | lo.mid | hi.mid | lo | hi |
| --- | --- | --- | --- | --- | --- | --- | --- |
| 0 | naive | $R_0$ | 12551 | 12505 | 12612 | 12414 | 12737 |
| 0 | naive | $\varepsilon_{2008}$ | 1.05 | 1.03 | 1.09 | 0.98 | 1.18 |
| 0 | naive | OFL<sub>2025</sub> | 919 | 911 | 933 | 892 | 956 |
| 0 | naive | OFL<sub>2034</sub> | 1139 | 1127 | 1151 | 1108 | 1174 |
| 3 | naive | $R_0$ | 12575 | 12517 | 12673 | 12411 | 12764 |
| 3 | naive | $\varepsilon_{2008}$ | 1.13 | 1.09 | 1.22 | 1.04 | 1.28 |
| 3 | naive | OFL<sub>2025</sub> | 919 | 906 | 943 | 887 | 959 |
| 3 | naive | OFL<sub>2034</sub> | 1151 | 1141 | 1167 | 1117 | 1185 |
| 3 | joint | $R_0$ | 12537 | 12484 | 12623 | 12340 | 12719 |
| 3 | joint | $\varepsilon_{2008}$ | 1.09 | 1.05 | 1.12 | 1.02 | 1.22 |
| 3 | joint | OFL<sub>2025</sub> | 912 | 903 | 935 | 875 | 953 |
| 3 | joint | OFL<sub>2034</sub> | 1141 | 1127 | 1156 | 1099 | 1170 |
| 5 | naive | $R_0$ | 12624 | 12558 | 12689 | 12461 | 12756 |
| 5 | naive | $\varepsilon_{2008}$ | 1.17 | 1.1 | 1.22 | 1.04 | 1.27 |
| 5 | naive | OFL<sub>2025</sub> | 927 | 913 | 939 | 896 | 960 |
| 5 | naive | OFL<sub>2034</sub> | 1160 | 1151 | 1172 | 1128 | 1182 |
| 5 | joint | $R_0$ | 12523 | 12484 | 12620 | 12361 | 12706 |
| 5 | joint | $\varepsilon_{2008}$ | 1.08 | 1.05 | 1.15 | 1 | 1.22 |
| 5 | joint | OFL<sub>2025</sub> | 911 | 899 | 929 | 878 | 953 |
| 5 | joint | OFL<sub>2034</sub> | 1140 | 1129 | 1153 | 1103 | 1164 |

**Figure S1.**
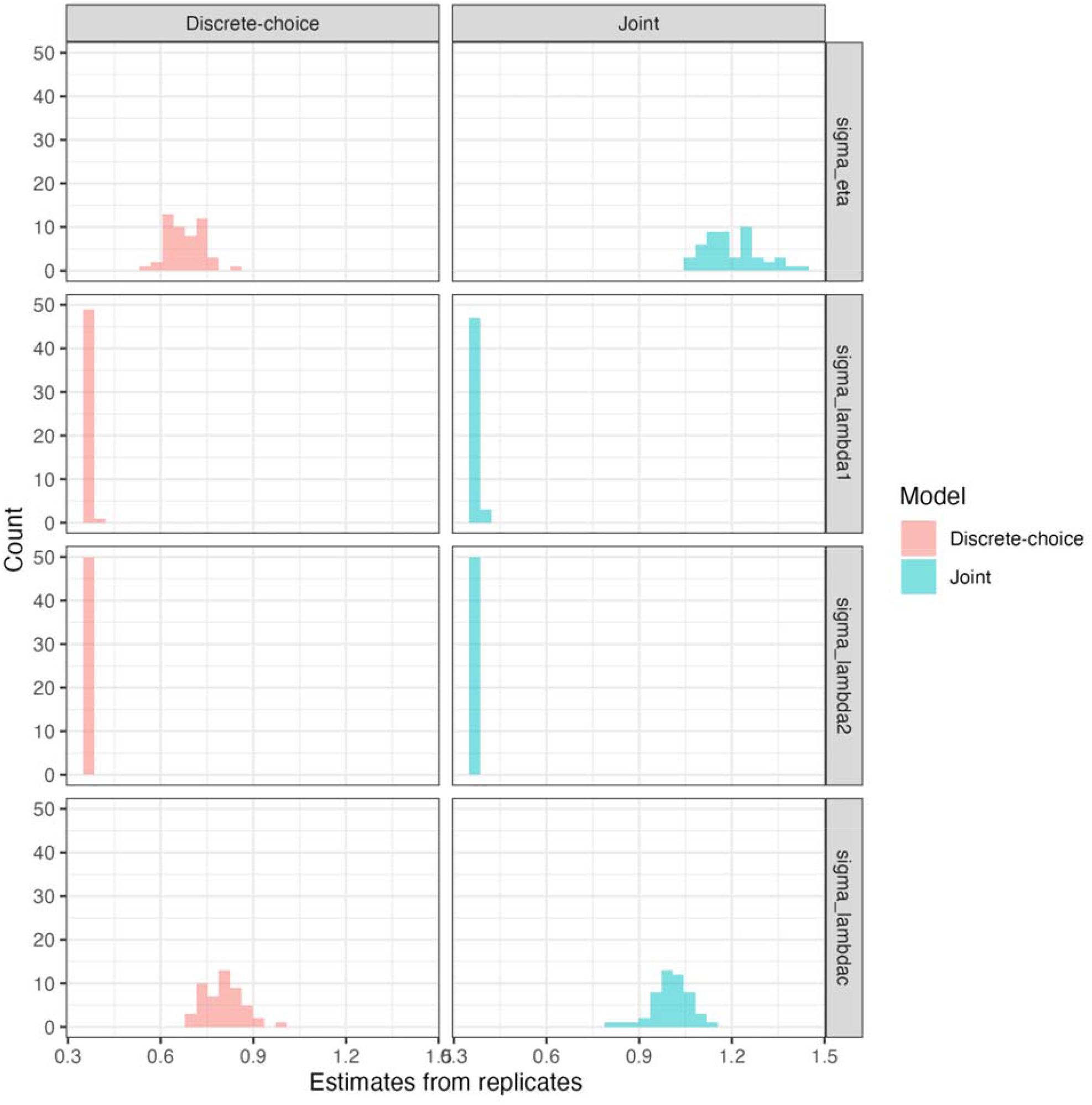
Estimated maximum likelihood estimates of variance parameters (rows) across data-generating mechanisms (columns) for parameters included in the discrete-choice estimation model. Each facet includes 25 replicate simulations, and parameter estimates that are concentrated in a narrow range indicate identifiability problems. A description of parameters and priors is in Table S1.

